# A sorghum Practical Haplotype Graph facilitates genome-wide imputation and cost-effective genomic prediction

**DOI:** 10.1101/775221

**Authors:** Sarah E. Jensen, Jean Rigaud Charles, Kebede Muleta, Peter Bradbury, Terry Casstevens, Santosh P. Deshpande, Michael A. Gore, Rajeev Gupta, Daniel C. Ilut, Lynn Johnson, Roberto Lozano, Zachary Miller, Punna Ramu, Abhishek Rathore, M. Cinta Romay, Hari D. Upadhyaya, Rajeev Varshney, Geoffrey P. Morris, Gael Pressoir, Edward S. Buckler, Guillaume P. Ramstein

**Affiliations:** Plant Breeding and Genetics Section, School of Integrative Plant Science, Cornell University, Ithaca, NY 14853; Chibas and Department of Agriculture and Environmental Sciences, Quisqueya University, Port-au-Prince, Haiti; Department of Agronomy, Kansas State University, Manhattan, Kansas; Institute for Genomic Diversity, Cornell University, Ithaca, NY 14853; International Crops Research Institute for the Semi-Arid Tropics (ICRISAT), Patancheru 502324, Telangana, India; United States Department of Agriculture-Agricultural Research Service, Robert W. Holley Center for Agriculture and Health, Ithaca, NY 14853

## Abstract

Successful management and utilization of increasingly large genomic datasets is essential for breeding programs to increase genetic gain and accelerate cultivar development. To help with data management and storage, we developed a sorghum Practical Haplotype Graph (PHG) pangenome database that stores all identified haplotypes and variant information for a given set of individuals. We developed two PHGs in sorghum, one with 24 individuals and another with 398 individuals, that reflect the diversity across genic regions of the sorghum genome. 24 founders of the Chibas sorghum breeding program were sequenced at low coverage (0.01x) and processed through the PHG to identify genome-wide variants. The PHG called SNPs with only 5.9% error at 0.01x coverage - only 3% lower than its accuracy when calling SNPs from 8x coverage sequence. Additionally, 207 progeny from the Chibas genomic selection (GS) training population were sequenced and processed through the PHG. Missing genotypes in the progeny were imputed from the parental haplotypes available in the PHG and used for genomic prediction. Mean prediction accuracies with PHG SNP calls range from 0.57-0.73 for different traits, and are similar to prediction accuracies obtained with genotyping-by-sequencing (GBS) or markers from sequencing targeted amplicons (rhAmpSeq). This study provides a proof of concept for using a sorghum PHG to call and impute SNPs from low-coverage sequence data and also shows that the PHG can unify genotype calls from different sequencing platforms. By reducing the amount of input sequence needed, the PHG has the potential to decrease the cost of genotyping for genomic selection, making GS more feasible and facilitating larger breeding populations that can capture maximum recombination. Our results demonstrate that the PHG is a useful research and breeding tool that can maintain variant information from a diverse group of taxa, store sequence data in a condensed but readily accessible format, unify genotypes from different genotyping methods, and provide a cost-effective option for genomic selection for any species.

## Introduction

The goal of plant breeding is to develop improved cultivars with high yield potential and better quality traits while reducing input requirements and environmental impact. Over time, plant breeding techniques have shifted from unconscious selection for domestication traits to deliberate selection and breeding program design. As the field of genetics developed, plant breeders began to use marker-assisted selection to associate genetic markers with desirable traits and inform breeding decisions (reviewed in Ramstein et al., 2018). First proposed by Meuwissen et al. in 2001, genomic selection (GS) is an extension of marker-assisted selection that uses genome-wide markers to make predictions about individual performance. Many studies have shown that genomic selection can accelerate the breeding process and rate of genetic gain without significantly increasing program costs (for example: Meuwissen et al., 2001; Bernardo and Yu, 2007; Heffner et al., 2010; Poland et al., 2012; Heslot et al., 2015; Muleta et al., 2019a). Increasingly dense marker or haplotype maps in major crops like maize (Bukowski et al., 2018) and sorghum (Lozano et al., 2019) can now be leveraged to inform breeding decisions.

Despite its apparent success, wide-scale adoption of genomic selection has been slow, particularly in developing countries where phenotypic selection is still cheaper than GS for many traits (Ribaut et al., 2010). GS costs depend on multiple components, including sample collection, processing, and sequencing costs, as well as bioinformatics requirements. Wider adoption of GS will require fast genotyping methods that are cheaper than phenotypic selection for all or most traits of interest. One way to make genotyping cheaper is to capitalize on pre-existing genomic data. Pan-genomic analyses capture a wider range of genetic variation and are becoming more common as genotyping costs decrease and the quantity of sequence data increases (for example: Alonso-Blanco et al., 2016; Morris et al., 2013; Shakoor et al., 2014). Tools that can extrapolate from existing whole-genome sequences (WGS) or other marker technologies and impute genotypes for individuals in a breeding population can utilize pre-existing WGS to minimize the amount of new data that breeders need to generate.

Here we introduce a method for genotyping breeding populations from low-coverage sequence data called the Practical Haplotype Graph (PHG). The PHG uses a graph of haplotypes to represent the variability in a breeding program and can merge genotypes from WGS and marker technologies. It takes a pan-genome approach to marker identification and relies on having a limited number of recombination events in each generation (Mace et al., 2009; Bouchet et al., 2017). In a breeding population, all major haplotypes in the progeny population are contained within a set of founder haplotypes. Moderate recombination shuffles founder haplotypes in subsequent generations, but progeny genotypes can be reconstructed if the approximate recombination positions can be identified. The PHG is designed to reconstruct haplotypes for a breeding population based on low-coverage random sequence, which reduces the quantity of new sequence data that needs to be collected for each cycle of GS.

In building a PHG database, moderate-to high-coverage WGS reads are aligned to the reference genome (Figure 1A) and collapsed into common, consensus haplotypes based on sequence divergence (Figure 1B, C). These consensus haplotypes are important for the PHG because they reduce the complexity of the graph, make it possible to associate traits with shared haplotypes, and fill in (impute) missing sequence information, while maintaining unique haplotypes. Once created, consensus haplotypes can be used to predict genotypes for new individuals. Skim sequence data are aligned to consensus haplotypes to find the best path through the graph, and SNP variants from the predicted haplotype path can be written to a VCF file. The result is a set of genome-wide SNP variant calls for each taxon, imputed from skim sequence (Figure 1D).

**Figure 1:**
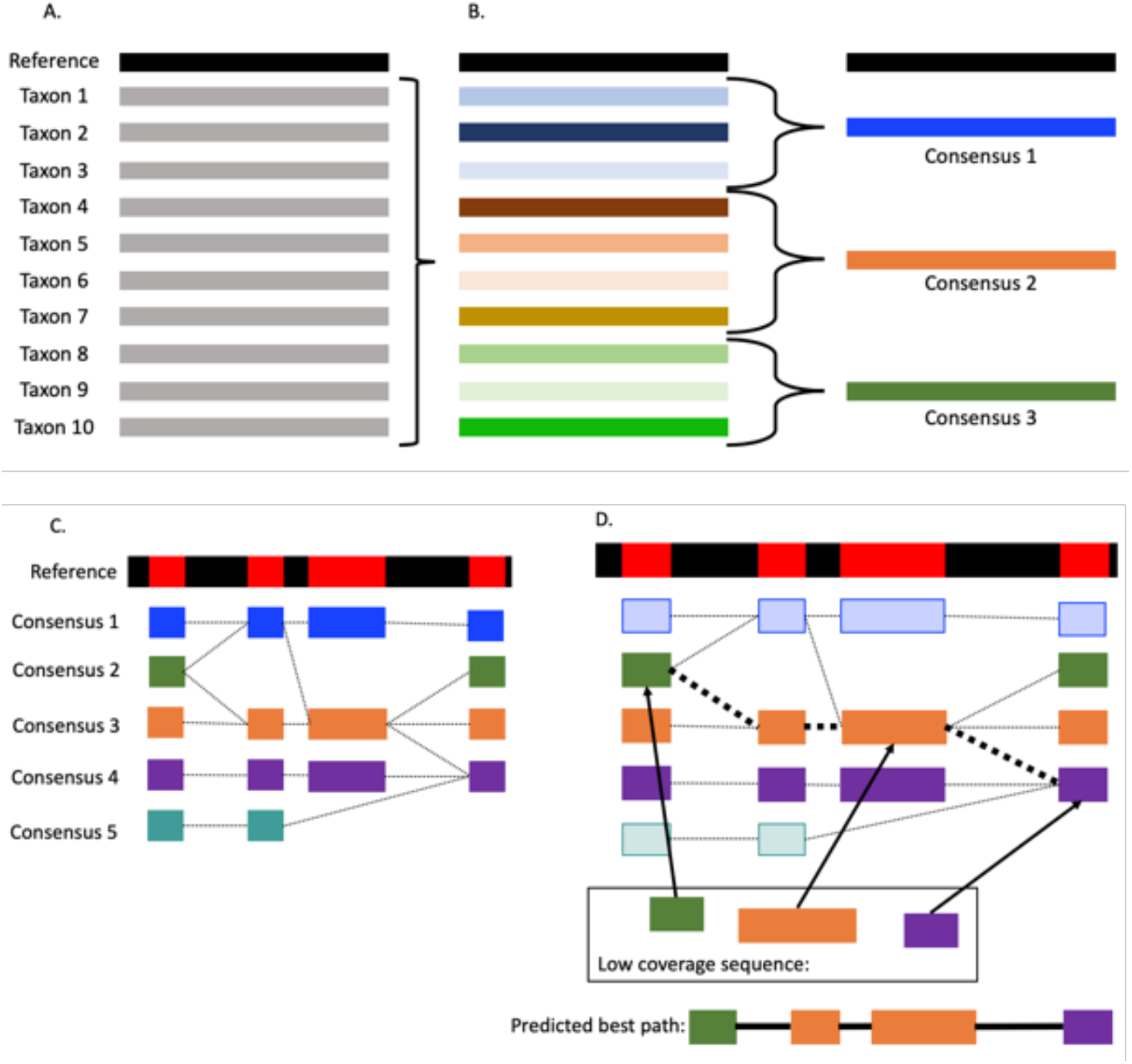
PHG database and haplotype creation. (A) WGS data are aligned to a reference genome and loaded into the PHG database. (B) A set of designated reference ranges is chosen and input data are condensed to produce consensus haplotypes at each reference range. Colors are used to indicate sequence similarity across taxa and only a single reference range is shown. (C) Unique consensus haplotypes are built for each reference range across the genome. Reference ranges are indicated as red regions of the black reference genome bar. (D) Low-coverage sequence data are aligned to the consensus haplotypes and a hidden Markov model links reference ranges across the genome to predict genome-wide haplotypes.

We tested the efficiency and accuracy of the PHG in a sorghum breeding program at Chibas in Port au Prince, Haiti. The climate in Haiti offers year-round growing conditions with three growing seasons annually. Currently, the primary sorghum breeding program at Chibas uses phenotypic recurrent selection and takes about a year per selection cycle. The main targets of selection include high yield, stalk sugar content, and sugarcane aphid resistance. GS has the potential to speed up the breeding cycle so that selections can be made up to three times per year, but is only useful if the potential increased genetic gain per cycle outweighs additional genotyping costs (Muleta et al., 2018). We tested a PHG built with 24 founders of the Chibas breeding program (referred to here as the “founder PHG”), and another built with the 24 Chibas founders plus an additional 374 taxa that are representative of overall sorghum diversity (referred to as the “diversity PHG”). With the Chibas breeding program as our test case, we evaluated whether the PHG can impute from low-coverage sequence, how PHG-imputed markers compare to other genotyping methods, and how much background genomic information is needed for a breeding program. Our results show that the PHG is an effective tool to impute genotypes for genomic selection from random skim sequence.

## Materials and Methods

### Phenotypic data

Phenotypes for height, brix, juice weight, leaf weight, earliness, stem weight, and grain yield were collected at the Regional Biofuels Technical and Knowledge Center field sites in Port au Prince, Haiti. The Chibas training population (250 taxa) was assayed in 5 experimental conditions (experiments) in Port au Prince, Haiti. Experiments differed by sowing date: (1) 9/7/2017, (2) 11/7/2017 to 11/10/2017, (3) 11/8/2017 to 11/10/2017 (irrigated), (4) 2/3/2018, and (5) 2/16/2018. All experiments were rainfed except for (3). Taxa were evaluated under randomized complete block designs with 4 replicates in each experiment, except for (4) and (5), which had 2 replicates each. For each trait, genotype (taxon) means were estimated as fixed effects under a linear mixed model with independent and identically distributed effects for experiments, interactions between genotypes and experiments, and replicates within experiments. Linear mixed models were fit with the R package lme4 (Bates et al. 2015).

### Sample collection and DNA extraction

Strips of mature leaf tissue measuring 4cm x 0.5cm were collected from month-old sorghum plants growing at the Chibas field site in Port au Prince, Haiti. Fresh tissue samples were placed in 96-well costar tube plates and lyophilized for 48 hours before being shipped to Intertek for DNA extraction. Samples were packed on silica beads and shipped at room temperature to the Intertek facility in Alnarp, Sweden, and DNA was extracted using their metal bead DNA extraction protocol.

### Library preparation and sequencing

DNA samples from 24 population founders were used to make TruSeq Nextera sequencing libraries at the Genomics facility at Cornell University. Samples from all 24 founders were pooled and sequenced in a single lane of 2 ×150 bp reads on an Illumina NextSeq500 instrument resulting in an average of 8x coverage per individual. The Chibas training population consists of 238 individuals. Samples in the training set were pooled in a single lane with 2,736 other individuals and sequenced at 2×150 bp reads on an Illumina NextSeq500 instrument, resulting in approximately 0.1x coverage for each individual. GBS data for comparison with PHG genotypes were from Muleta et al. (2019b).

### Building the sorghum PHG

A sorghum practical haplotype graph was built using scripts in the p_sorghumphg bitbucket repository and PHG version 0.0.9. Instructions for building a new PHG can be found on the PHG Wiki, available on Bitbucket at https://bitbucket.org/bucklerlab/practicalhaplotypegraph/wiki/Home (Figure 2).

**Figure 2:**
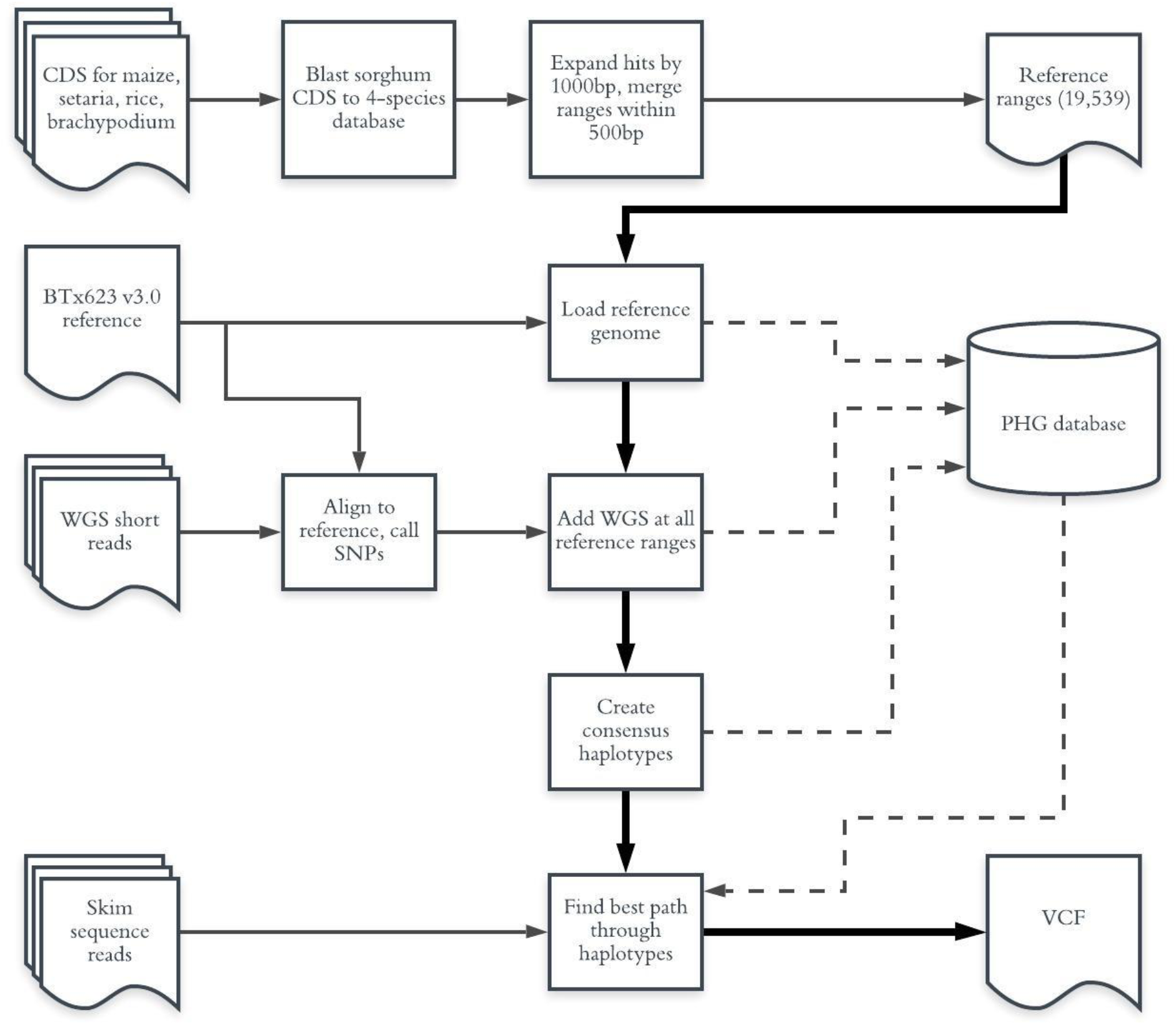
Steps in building and using the sorghum PHG. The main PHG processing steps are indicated with bold arrows. Steps that store or read information from the PHG database are indicated with dotted lines.

#### Creating and loading reference ranges

Reference ranges for the PHG were chosen based on conserved gene annotations. Coding sequences (CDS) from the sorghum version 3.1 genome annotations and the version 3.0 reference genome were downloaded from JGI and compared to a BLAST database containing CDS for *Zea mays, Setaria italica, Brachypodium distachyon*, and *Oryza sativa* (Schnable et al., 2009; Bennetzen et al., 2012; Vogel et al., 2010; Ouyang et al., 2007) that was created using BLAST+ command line tools (Altschul et al., 1997). The sorghum version 3.1 CDS annotations and version 3.0 reference genome (McCormick et al., 2017) were compared to the 4-species database with blastn default parameters. Sorghum gene intervals were kept if there was at least one hit to the four-species database, and gene start and end coordinates were used to create initial reference intervals. Initial gene intervals were expanded by 1000 bp on either side of the gene coordinates, and intervals within 500 bp of each other were merged to form a single reference range. The resulting dataset contains 19,539 intervals spaced across the genome, which we designated “genic reference ranges”, while the intervals between genic reference ranges were added to the database as 19,548 “intergenic reference ranges.” The LoadGenomeIntervals pipeline was used to add reference genome sequence to the database for both genic and intergenic ranges, while sequence data from additional taxa were added only to the genic reference ranges.

#### Adding haplotypes from diverse taxa and creating consensus haplotypes

Sequence data were aligned to the version 3.0 sorghum BTx623 reference genome with BWA MEM (McCormick et al., 2017; Li & Durbin, 2009). Taxa in the PHG are as follows: 24 founder individuals from the Chibas sorghum breeding program, 274 previously-published taxa (42 from Mace et al., 2013, 232 from Valluru et al., 2019), and 100 taxa from the ICRISAT mini-core collection, for a total of 398 taxa. No *de novo* genome assemblies are included. Variants were called with Sentieon’s HaplotypeCaller pipeline (Sentieon DNAseq, 2018) and the resulting gVCF files were added to the PHG using the CreateHaplotypesFromGVCF pipeline. The same process was used to make a smaller PHG database with only the 24 founder individuals from the Chibas breeding program.

The 398 or 24 individuals added to the PHG were condensed into common “consensus haplotypes” using the CreateConsensus pipeline. Taxa were clustered with the UPGMA clustering method in TASSEL (Bradbury et al., 2007) at ten possible divergence levels: 0.001, 0.00075, 0.0005, 0.00025, 0.0001, 0.000075, 0.00005, 0.000025, 0.00001, and 0. Unique divergent haplotypes were retained as consensus haplotypes (minTaxa=1), and all other parameters were left at their default values. The Chibas founder taxa were used as a baseline to select the best level of haplotype collapse for this dataset, and a consensus level of 0.00025 (1 in 4000 bp differences) was chosen. This level maximizes SNP count and concordance with other sequencing methods, while minimizing the number of haplotypes at each reference range (Figure 3). Subsequent genotyping and imputation analyses were run using this consensus level.

**Figure 3:**
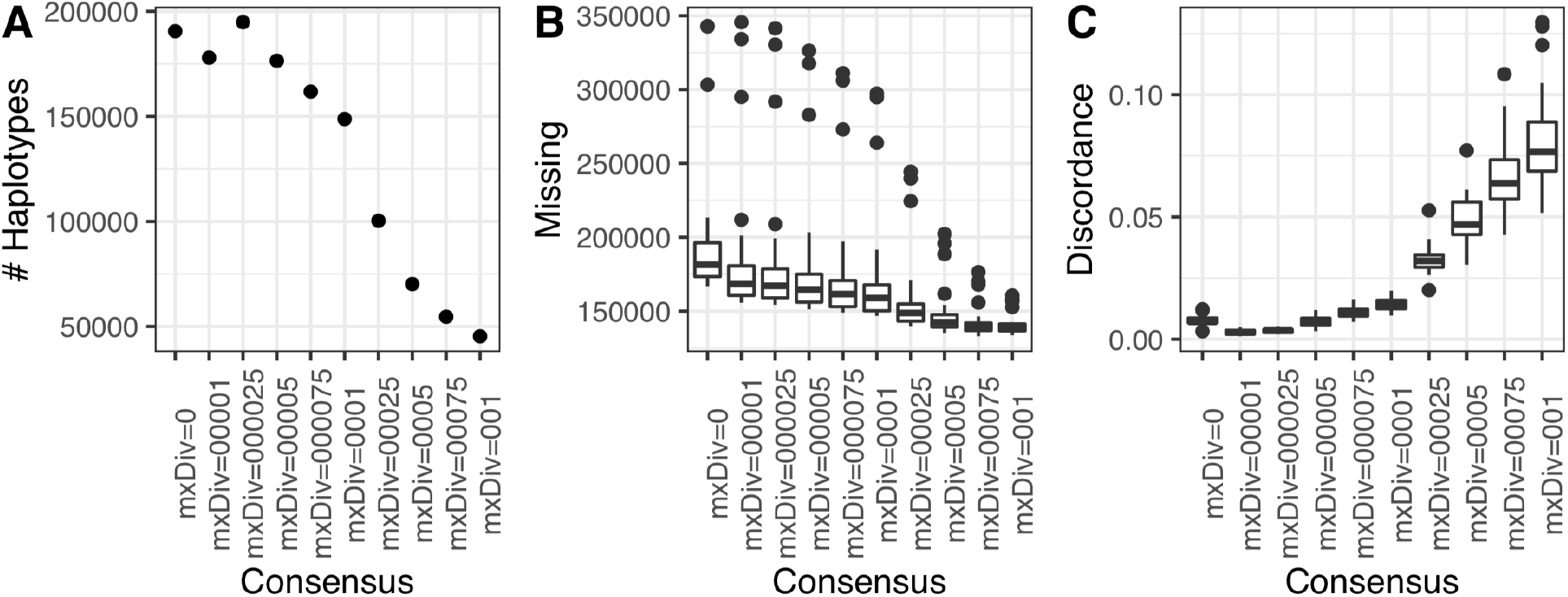
Metrics used to choose a consensus level for condensing haplotypes in the founder PHG. A) Number of haplotypes and B) Missing counts at each consensus level. C) Discordance between SNPs called by the PHG and a set high-confidence WGS SNPs. The divergence level of mxDiv=0.00025 is at an inflection point in A) and was chosen as the level that balances the tradeoff between amount of missing information in the database and discordance from collapsing haplotypes.

#### Predicting new genotypes

After creating consensus haplotypes, the findPaths pipeline maps reads to the consensus pangenome and uses a hidden Markov model (HMM) and the Viterbi algorithm (Rabiner, 1989) to predict the best path through all consensus haplotypes in the graph. In each analysis we subset the pangenome graph to only the consensus haplotypes containing one or more of the 24 taxa that founded the Chibas breeding program. Paired-end 150 bp reads with varied sequence coverage from random sequence data were used as input to predict the best path through the graph, both for individuals in the database and for new taxa that are not already represented. Paths were exported to a VCF file and used to evaluate accuracy of imputation and genomic prediction with the PHG.

All steps described above were run using a docker image containing necessary software and scripts available for download from the PHG Bitbucket site. Specific parameters used to run each script can be found in the config.txt file on bitbucket. Additional scripts and files for creating the sorghum PHG are available at https://bitbucket.org/bucklerlab/p_sorghumphg/src/master.

### Imputation Accuracy

#### PHG imputation accuracy for WGS

WGS data for the Chibas founder taxa were down-sampled with seqtk (Li, 2013) to 1x, 0.1x, and 0.01x coverage. Sequences were produced with three separate seed integers to create three unique sets of reads at each level of coverage. The full WGS data and each set of down-sampled sequencing reads were run through the PHG findPaths pipeline using a PHG database with nodes built from the Chibas founders, minReads=0, minTaxa=1, and all other parameters left at default values. Setting the minReads parameter to 0 means that the HMM will attempt to find a path through the entire genome, even when there is no sequence data observed at a particular reference range. Setting the minTaxa parameter to 1 means that all haplotypes are kept, even if taxa are too divergent to group with other individuals in the database. SNPs were written at all variant sites in the graph, as well as all positions in the sorghum hapmap (Lozano et al, 2019). SNP calling accuracy was assessed by comparing PHG SNP calls to a set of 3,468 GBS SNPs (Muleta et al., 2019b). SNPs with minor allele frequency < 0.05 or call rate < 0.8 were removed before comparing PHG and GBS SNP calls. Haplotype calling accuracy was evaluated by running low-coverage taxa through the database and counting the number of times that the selected node in the graph contained the taxon being imputed.

While error rates for most taxa were consistent with the overall error, BF-95-11-195 stood out as having a 5-fold higher error than expected in calling SNPs, although its haplotype calling error was not abnormally high. We suspect this sample was mixed up or contaminated with DNA from another sample during sequencing, but kept BF-95-11-195 in the database and included it in all analyses.

#### Beagle 5.0 imputation accuracy

Because the PHG is expected to be useful when only skim sequence data is available for an individual, we compared PHG imputation accuracy to Beagle 5.0 (Browning & Browning, 2016) imputation accuracy from low-coverage sequence. WGS data for each taxon was down-sampled as described above. Each down-sampled dataset and the full-coverage (∼8x) WGS data from 24 founders of the Chibas sorghum breeding program was aligned to the sorghum v3.0 reference genome with BWA MEM (McCormick et al., 2017; Li & Durbin, 2009) and variants were called with the Sentieon DNASeq variant calling pipeline (Sentieon DNAseq, 2018). VCF files for each founder were merged using bcftools (Li et al., 2009). When variant sites did not line up in the full coverage WGS (i.e., a variant was called for one individual but not for another such that merging variant calls across taxa would produce a missing call in some taxa and an alternate allele call in others), the unobserved site was assumed to be the reference call. To simplify both the Beagle and PHG imputation pipelines and because individuals used in the database construction were expected to be inbred lines, all heterozygous calls were assumed to come from sequencing and genotyping errors rather than residual heterozygosity, and were removed. For the downsampled datasets, unobserved sites were left as missing. A reference panel created from full-coverage WGS was used to impute SNPs in the down-sampled VCF files. No sites in the downsampled data were masked; instead, missing information was imputed directly using the reference panel. In the full-coverage dataset, 1% of all sites were masked and re-imputed. Imputation accuracy at all levels of sequence coverage was evaluated by comparing Beagle calls to a set of 3,849 GBS SNPs.

#### Designing amplicon sequencing markers and using the PHG to impute variants

Amplicon sequencing technologies like rhAmpSeq use PCR amplification to identify SNPs at targeted sites in the genome (Fresnedo-Ramírez et al., 2019). GS with custom rhAmpSeq markers was tested with and without PHG imputation. These rhAmpSeq markers were developed using 100 taxa from the ICRISAT mini-core collection, which are also used in the diversity PHG (Supplemental Table 1). Paired SNP variants between 10-100 bp apart were identified in this panel of 100 taxa and designated as potential haplotype regions. Each potential haplotype region was expanded on either side of the SNP pair to generate 104bp segments centered on the initial pair of SNPs. This identified 336,082 potential haplotype regions, and the polymorphic information content (PIC) score was calculated for each haplotype using the 100-taxa panel.

The sorghum reference genome annotation (Sbicolor 313, annotation v3.1) and sequence (Sbicolor 312, assembly v3.0) were used to divide the chromosome-level assembly into 2904 genomic regions. Each region contained equal numbers of non-overlapping gene models; overlapping gene models were collapsed into a single gene model. 2892 of these regions contained at least one SNP-pair haplotype. For each region, the SNP-pair haplotype with the highest PIC score was selected as a representative marker locus. These genome-wide candidates, along with 148 target marker regions of interest provided by the sorghum breeding community, were used by the rhAmpSeq team at Integrated DNA Technologies, Inc. to design and test rhAmpSeq genotyping markers. After design and testing, markers for 1974 genome-wide haplotype targets and 138 community-identified targets were selected as the rhAmpSeq amplicon set.

The rhAmpSeq sequence data was processed through the PHG findPaths pipeline in the same way as the random skim sequence data described above. To determine how many markers the PHG needs for imputation, we randomly sampled 500 and 1000 loci from the original set of 2,112 haplotype targets and used the PHG findPaths pipeline to impute SNPs across the rest of the genome. Results were written to a VCF file and used for genomic prediction.

### Genomic prediction

PHG SNP performance in genomic prediction was evaluated using a set of 207 individuals in the Chibas training population for which GBS (Elshire et al., 2011) and rhAmpSeq SNP data was also available. PHG genotypes were predicted with the findPaths pipeline of the PHG using either random skim sequence data at approximately 0.1x or 0.01x coverage, or rhAmpSeq reads for 2112, 1000, or 500 loci (corresponding to 4854, 1453, and 700 SNPs, respectively) as inputs. Paths were determined by using an HMM to extrapolate across all reference ranges (minReads=0, removeEqual=false). Genomic relationship matrices based on PHG-imputed SNPs were created with the “EIGMIX” option in the SNPRelate R package (Zheng et al., 2012). A haplotype relationship matrix using PHG consensus haplotype IDs was created as described in Jiang, Schmidt, & Reif (2018) equation (2), using the tcrossprod function in base R. For GBS markers, markers with more than 80% missing or minor allele frequency <= 0.05 were removed from the dataset and missing markers were imputed with mean imputation, and a genomic relationship matrix was computed as described in Endelman et al. (2011). Genomic prediction accuracies were Pearson’s correlation coefficients between observed and predicted genotype means, calculated with 10 iterations of 5-fold cross validation. GBS and rhAmpSeq SNP data without PHG imputation were used as a baseline to determine prediction accuracy. To see if the PHG could impute WGS starting from rhAmpSeq amplicons, genomic prediction accuracies using the PHG with rhAmpSeq-targeted loci were compared to prediction accuracies using rhAmpSeq data alone.

## Results

We developed two sorghum PHG databases. One contains only the original founder haplotypes of the Chibas breeding population (“founder PHG”, 24 genotypes), while the other PHG contains both the Chibas founders and WGS from the founder taxa plus an additional 374 taxa that reflect the overall diversity within sorghum (“diversity PHG”, 398 genotypes). We determined how much sequence coverage is needed for the PHG and how genomic prediction with PHG-imputed markers compares to genomic prediction with GBS and rhAmpSeq markers. Data was processed through the founder PHG and the diversity PHG in the same way.

### A PHG for global sorghum diversity

The sorghum diversity PHG stores sequence information for 398 diverse inbred lines at 19,539 reference ranges covering all genic regions of the genome, and is built from WGS data with coverage ranging from 4 to 40x, although most individuals have 10x coverage or less. The founder PHG contains WGS at ∼8x coverage for 24 founders of the Chibas breeding program. A gVCF file is made by calling variants between WGS and the reference genome, and variants from the gVCF are added to the PHG database in all genic reference ranges. The reference ranges in both versions of the sorghum PHG are centered around gene regions. At each reference range, haplotypes are collapsed into consensus haplotypes to combine similar taxa and fill in missing sequence across the graph. There is a tradeoff when choosing a divergence cutoff for consensus haplotypes: a low divergence level will retain lower-frequency SNPs, but not fill in gaps and missing data as well as a high divergence level. In both the diversity PHG and the founder PHG, consensus haplotypes were created by collapsing haplotypes that had fewer than 1 in 4,000 bp differences (mxDiv=0.00025), which is a slightly lower density of variants than the GBS SNP density reported by Morris et al. (2013). This level was chosen because it marks an inflection point in the number of consensus haplotypes that are created (Figure 2A), with an average of 5 haplotypes per reference range in the founder PHG and intermediate levels of missingness and discordance with WGS calls made with the Sentieon pipeline (Figure 2B-C). The consensus haplotypes produced at this divergence level were used to evaluate PHG SNP-calling and genomic prediction accuracy.

### PHG SNP-calling accuracy is minimally affected by read count

PHG haplotype and SNP calling accuracies are minimally affected by decreasing amounts of sequence data. The PHG was evaluated to determine the lower boundary of sequence coverage before imputation accuracy decreased substantially. For each founder in the Chibas breeding program, WGS was subset down to ∼2.4M, 243,333, and 24,333 reads, corresponding to 1x, 0.1x, and 0.01x genome coverage, respectively. Sequencing reads were randomly selected from the original WGS fastq files and used to predict SNPs or haplotypes with the PHG, and PHG-predicted SNPs and haplotypes at each level of sequence coverage were evaluated for accuracy. Haplotypes were considered correct if the imputed haplotype node for a given taxon also contained that taxon in the PHG. SNPs were considered correct if they matched GBS calls at 3,369 loci for which GBS data had MAF > 0.05 and a call rate > 0.8.

Haplotype error was higher than SNP calling error in both the founder PHG database (24 taxa) and the diversity PHG database (398 taxa), and accuracy increased in both databases with increasing sequence coverage. Haplotype error ranged from 11.5-12.1% error in the founder database, to 18.6-23.5% error in the diversity database. SNP error ranged from 2.9-5.9% and 4.3-15.2% in the founder and diversity PHG databases, respectively (Figure 4). Higher haplotype error rates are likely due to similarity among haplotypes that leads the HMM to call an incorrect haplotype even if most of the SNPs within that haplotype are correct.

**Figure 4:**
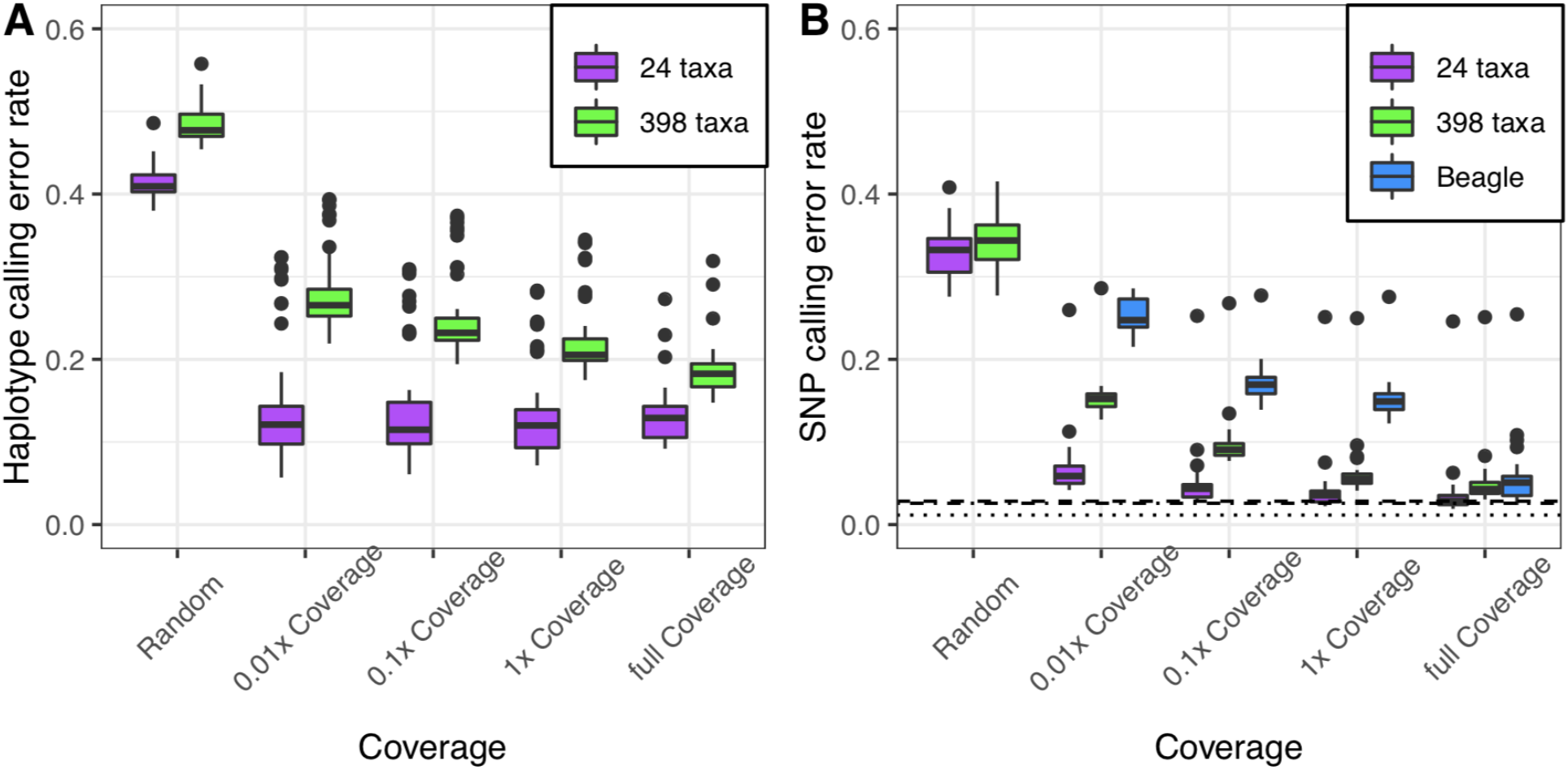
PHG haplotype (A) and SNP (B) error rates compared to GBS data with the 24-taxa founder PHG database (purple) and the 398-taxa diversity PHG database (green) for a random path through the PHG and a range of sequence coverages. Beagle SNP calling accuracy at each coverage level is included in B (blue). Horizontal lines in B represent baseline error rate of the PHG (dotted), GBS (dot-dash), and Beagle (dashed).

PHG-imputed SNP calls were better than Beagle-imputed SNP calls when using low-coverage skim sequence, and about equal to Beagle calls at full coverage (∼8x) WGS. Notably, while SNP calling accuracy did improve with higher sequence coverage for all imputation methods, calling SNPs with the founder PHG resulted in only a 3% difference in SNP-calling accuracy between the highest coverage tested (8x coverage, 2.9% error) and the lowest coverage tested (0.01x coverage, 5.9% error; Figure 4B). In contrast, there was a 23% difference in SNP-calling accuracy between low and high coverage sequence when using Beagle for imputation. These results indicate that the PHG is a useful imputation tool that can impute SNPs more accurately than Beagle when starting from low-coverage skim sequence.

To determine the PHG baseline error rate, we looked at the intersection of PHG, Beagle, and GBS SNP calls at 3,363 loci in 24 taxa. These loci were chosen because they represented biallelic SNPs called with the GBS pipeline that also had genotype calls made by both the PHG and Beagle imputation methods. The baseline error was calculated as the proportion of SNPs where genotype calls from one of the three methods did not match the other two. Using this metric, baseline error for Beagle imputation, GBS SNP calls, and PHG imputation were calculated to be 2.83%, 2.58%, and 1.15%, respectively (Figure 3B, dashed and dotted lines). To investigate the source of the 1.15% PHG error, we compared the SNP calls from a model path through the PHG (i.e., the calls that the PHG would make if it called the correct haplotype for every taxon at every reference range) to the incorrect PHG SNP calls. Allele calls that were correct in the model SNP set but not called in the genotypes predicted by the findPaths pipeline were counted as an error in the pathfinding step, which is caused by the HMM incorrectly calling the haplotype at a reference range. Allele calls that were not present in the model SNP set were counted as an error in the consensus step. Consensus errors are due to alleles being merged in the createConsensus pipeline because of similarity in haplotypes. Our analysis found that 25% of the PHG baseline error comes from incorrectly calling the haplotype at a given reference range (pathfinding error), while 75% comes from merging SNP calls when creating consensus haplotypes (consensus error). Haplotype and SNP calls from the founder PHG were more accurate than calls with the diversity PHG at all levels of sequence coverage. These results indicate that haplotype and SNP calls from the founder PHG are more accurate than those from the diversity PHG. Therefore, subsequent analyses were done with the founder PHG.

We compared accuracy in calling minor alleles between PHG and Beagle SNP calls. Beagle accuracy drops when dealing with datasets where 90-99% of sites are missing (0.1 or 0.01x coverage) because it makes more errors when calling minor alleles (Figure 5, red circles). When imputing from 0.01x coverage sequence, the PHG calls minor alleles correctly 73% of the time, whereas Beagle calls minor alleles correctly only 43% of the time. The difference between PHG and Beagle minor allele calling accuracy decreases as sequence coverage increases. At 8x sequence coverage, both methods perform similarly, with minor alleles being called correctly 90% of the time. PHG accuracy in calling minor alleles was consistent regardless of minor allele frequency (Figure 5, blue triangles).

**Figure 5:**
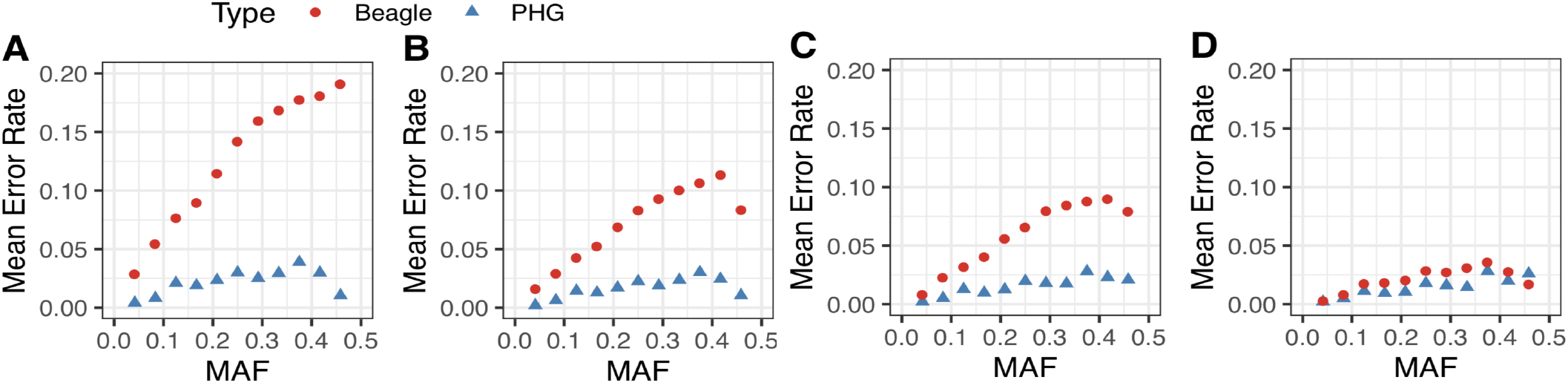
Mean error in calling minor alleles with Beagle (red circles) and the PHG (blue triangles). (A) 0.01x coverage, (B) 0.1x coverage, (C) 1x coverage, (D) full (8x) coverage.

### PHG genomic prediction accuracies match genomic prediction accuracies from GBS

To test whether PHG haplotype and SNP calls predicted from low-coverage sequence are accurate enough to use for genomic selection in a breeding program, we compared prediction accuracies with PHG-imputed data to prediction accuracies with GBS or rhAmpSeq markers. We predicted breeding values for 207 individuals from the Chibas training population for which GBS, rhAmpSeq, and random skim sequencing data was available. Haplotype IDs from PHG consensus haplotypes were also tested to evaluate prediction accuracy from haplotypes instead of SNPs (Jiang, Schmidt, & Reif, 2018). The 5-fold cross-validation results suggest that prediction accuracies for SNPs imputed with the PHG from random skim sequences are similar to prediction accuracies from GBS SNP data for multiple phenotypes, regardless of sequence coverage for the PHG input. Haplotypes can be used with equal success; prediction accuracies using PHG haplotype IDs were similar to prediction accuracies using PHG or GBS SNP markers (Figure 6A). Results are similar with the diversity PHG database (Supplemental Figure 1). With rhAmpSeq markers, adding PHG-imputed SNPs matched, but did not improve, prediction accuracies relative to accuracy with rhAmpSeq markers alone (Figure 6B). Using the PHG to impute from random low-coverage sequence can, therefore, produce genotype calls that are just as effective as GBS or rhAmpSeq marker data, and SNP and haplotype calls predicted with the findPaths pipeline and the PHG are accurate enough to use for genomic selection in a breeding program.

**Figure 6:**
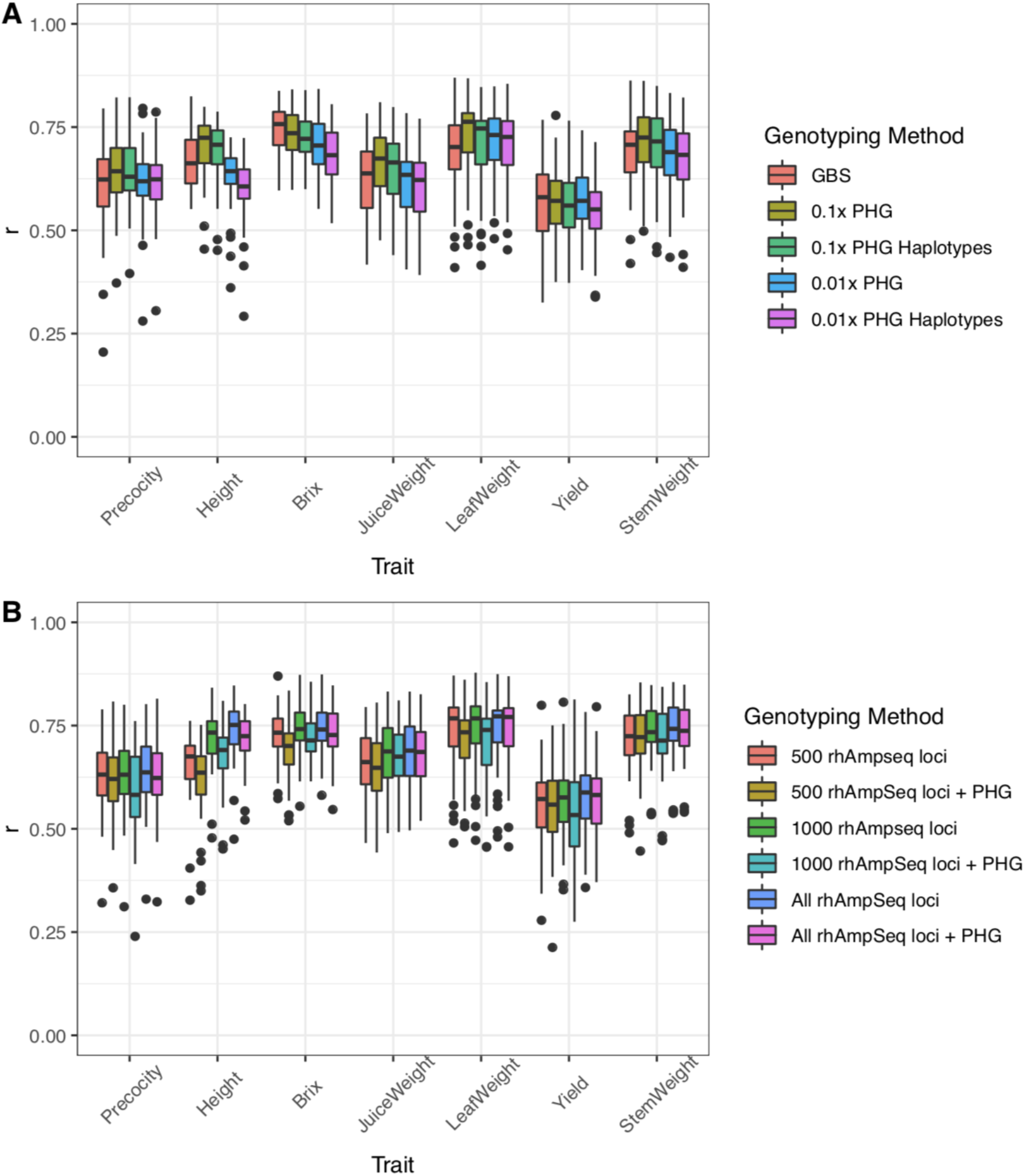
5-fold cross validation prediction accuracies for the Chibas training population (n=207, 10 iterations) are the same when using A) GBS, PHG SNPs with 0.1x and 0.01x sequence coverage, or PHG haplotypes. B) Prediction accuracies do not change when using rhAmpSeq markers alone or using rhAmpSeq markers with additional markers from the PHG, regardless of how many rhAmpSeq markers are included.

## Discussion

### SNP calling accuracy

The PHG is a cost-effective genotyping tool that combines WGS data in a database to capture the main haplotype groups in the breeding program or species. Sequences and consensus haplotypes stored in the PHG can be used for genomic prediction. We built a diversity PHG with 398 individuals to capture sorghum-wide diversity and a second, smaller database with only the 24 breeding program founders. In general, the 24-taxa founder PHG database had higher SNP and haplotype calling accuracy, but both databases produced genotypes that could be used effectively for genomic prediction.

When testing the accuracy of the PHG, we find that random skim sequence data can be imputed for SNPs across the PHG reference ranges with high accuracy. Based on the levels tested, 0.01x coverage is the most cost effective level of sequence coverage with 94.1% SNP calling accuracy - only a 3% drop in SNP calling accuracy relative to accuracy at 8x-coverage WGS. For the sorghum genome, 0.01x coverage corresponds to ∼25,000 completely random paired-end 150 bp reads. The sequence reads tested here were selected randomly and are unlikely to cover all reference ranges, which shows that the PHG can impute across reference ranges even when sequence can only be aligned to a portion of the ranges in the database. Long-read sequence data, which creates fewer reads, could, therefore, also be used as input for the PHG path-finding algorithm (findPaths pipeline). A few long reads spaced randomly across the genome would likely identify haplotypes with similar levels of accuracy as 0.01x coverage short-read sequence data.

### Genomic prediction accuracy

Both 0.01x and 0.1x coverage sequence imputed with the PHG, as well as haplotype IDs from the PHG, can be used for genomic prediction with prediction accuracies similar to those produced by GBS markers. In the training dataset comprising 207 individuals, there was no difference in using a haplotype relationship matrix instead of genomic relationship matrix built from PHG SNPs. However, in larger datasets with more individuals, using haplotype IDs instead of SNP markers may improve computational efficiency without a cost in terms of prediction accuracy. Using the PHG with rhAmpSeq markers worked as well as using rhAmpSeq markers alone for complex traits, but prediction accuracies dropped slightly for some traits (e.g. height, juice weight) if only 500 rhAmpSeq markers were used with PHG imputation. This could be related to trait genetic architecture; height is an oligogenic trait in sorghum, while traits like grain yield and precocity would be expected to be more polygenic (Pereira and Lee, 1995; Girma et al., 2019).

Low-coverage sequence in conjunction with the PHG allows large-scale imputation of progeny genotypes across the genome. We show that the PHG can be used with random skim sequence and rhAmpSeq sequences, suggesting that the PHG can also be used to unify sequence data from diverse genotyping platforms. With only 0.01x genome coverage needed for accurate calls, per-taxon sequencing costs become negligible compared to the cost of DNA extraction and library preparation (both of which are also continuing to decrease: see Anderson et al., 2018; Baym et al., 2015; Zou et al., 2017). These new technologies, together with the PHG, provide breeders with an effective, inexpensive, and fast tool for imputation and genomic selection that works across genotyping platforms. Since the founder PHG produces more accurate SNP and haplotype calls than the diversity PHG, we also suggest that small PHG databases built for specific breeding programs can be more effective than a single species-wide haplotype graph.

It is surprising that the diversity PHG had lower SNP and haplotype calling accuracy than the founder PHG because it contains more information about the species diversity overall. We hypothesize that the difference in performance is due to differences in allele frequency or LD patterns between the Chibas breeding program and the taxa in the diversity PHG. As expected for a breeding program, the Chibas breeding material captures much less diversity than is present in the species as a whole. If common alleles in the Chibas founder individuals are rare relative to the taxa included in the diversity PHG, then the Chibas founders may have been pulled into consensus haplotypes with alleles that are not common in the Chibas breeding program. Using these consensus haplotypes to impute from skim sequence could have added alleles not present in the founder PHG which would increase SNP calling error relative to GBS. Linkage disequilibrium patterns between sorghum races may also differ. Therefore, transition probabilities between reference ranges may be estimated less accurately in a diverse PHG database comprising multiple races. For now, if working in a breeding context where diversity is limited, a founder PHG with program-specific haplotypes appears to work best. In the future, this issue could be solved by anchoring consensus haplotypes to a specific set of taxa (in this case, the Chibas founders). Other taxa in the database would only be used to fill in missing information in the set of anchored consensus haplotypes. The resulting haplotypes would have less missing information than the current founder PHG, but would maintain the allele frequencies and haplotype patterns of the original set of anchor taxa.

### Decreasing genotyping costs

The cost of building a PHG depends on the number of individuals for which WGS or *de novo* assemblies need to be produced. Relying on existing resequencing data when possible can significantly reduce the overall cost. For the sorghum PHGs produced here, all taxa in the Chibas sorghum breeding program were multiplexed in a single sequencing lane, resulting in approximately 8x coverage for each individual and low levels of missing data. The initial sequencing investment for the founder PHG was $5,283. The additional 374 taxa added to the diversity PHG were produced for other research purposes and no additional sequence data were produced for these individuals. Thus, the total upfront cost for building the sorghum PHGs was under $6,000 - less than the genotyping costs for a round of genomic selection.

The PHG aims to make genotyping and genomic selection marker agnostic, i.e. all marker systems should produce similar results. We see the PHG likely to be used with five current and future platforms. The most expensive is GBS at approximately $15/sample, which is substantially driven by expensive DNA preparation and uneven library coverage. Tn5 based skim sequencing can use simple DNA extraction protocols, has the same procedure for any species, and costs ∼$10/sample. Targeted amplicon-based sequencing can use very inexpensive sample preparation protocols and provides 500 to 2000 loci for $3.50 to $10 per sample. It does, however, require significant upfront investment to develop amplicons for each species, and the per-sample cost is dependent on the number of samples processed annually. The cost of random prime sequencing with simple DNA extractions is similar to targeted amplicon sequencing at $5 to $10 per sample, but the price for random prime sequencing does not depend on sample number. Long-read sequence data can also be used with the PHG and the price is likely to drop as long-read technologies are developed further. The PHG is designed to work with any of these sequence types, making it possible to unify sequence data from across multiple platforms.

### Turnaround time

Continuous GS cycles require rapid data turnaround so that there is enough time to identify the best progeny and plan crosses before flowering. In the Chibas breeding program, tissue samples can be collected about 1 month after planting. For continuous GS cycles to be possible, all sample processing, sequencing, and variant calling must be complete 6 weeks after samples are collected. Using a DNA extraction service took two weeks, with another 1.5 weeks needed for library preparation and sequencing. Sequencing data took 48 hours to run through the founder PHG, giving us about a 4-week turnaround time from sample collection to genotype data and leaving two weeks to make selections and plan crosses. Additional time could potentially be saved with cheap, on-site DNA extractions and library preparations (Anderson et al., 2018; Zou et al., 2017). The extra time gained with on-site methods could make a second round of tissue collection and sequencing possible if samples are contaminated or if more data is needed for a particular breeding line. With these improvements, genotyping would no longer be the factor limiting turnaround time, and techniques like speed breeding could be used to further reduce cycle time in sorghum breeding (Watson et al. 2018).

### Conclusions

We developed two practical haplotype graphs: a diversity PHG with 398 individuals consisting of all 5 sorghum races and a founder PHG with individuals specific to the Chibas sorghum breeding program, which consists mostly of breeding lines derived from east African caudatum sorghums (Muleta et al., 2019b). Our study demonstrated how the PHG can accurately impute SNPs and haplotypes from low coverage random sequence data and predict genotypes for a breeding population. The PHG was able to produce better imputed genotype calls than Beagle 5.0 when using low-coverage random sequence data, and prediction accuracies from PHG-imputed SNPs matched those produced with higher-cost GBS marker data for traits with a range of genetic architectures when starting from similar levels of sequence coverage. These results demonstrate that the PHG is a useful low-cost genotyping tool for breeders around the globe.

## Acknowledgments

This study is made possible by the support of the American People provided to the Feed the Future Innovation Lab for Collaborative Research on Sorghum and Millet through the United States Agency for International Development (USAID) under Associate Award No. AID-OAA-LA-16-00003. The information, data, or work presented herein was also funded in part by the Advanced Research Projects Agency-Energy (ARPA-E), U.S. Department of Energy, under Award Number DE-AR0000598. The contents are the responsibility of the authors and do not necessarily state or reflect those of USAID, the United States Government, or any agency thereof. We thank Integrated DNA Technologies for a custom rhAmpSeq panel in sorghum. Additional support comes from the United States Department of Agriculture, Agricultural Research Service and the Bill and Melinda Gates Foundation.

## Data availability

Raw sequence data for this project is available at NCBI BioProject ID PRJNA575060. All processed data can be found on CyVerse at https://doi.org/10.25739/nccb-0k78. Scripts to replicate the analyses can be found on bitbucket at https://bitbucket.org/bucklerlab/p_sorghumphg/src/master/.

## Supplemental figure

**Supplemental Figure 1:**
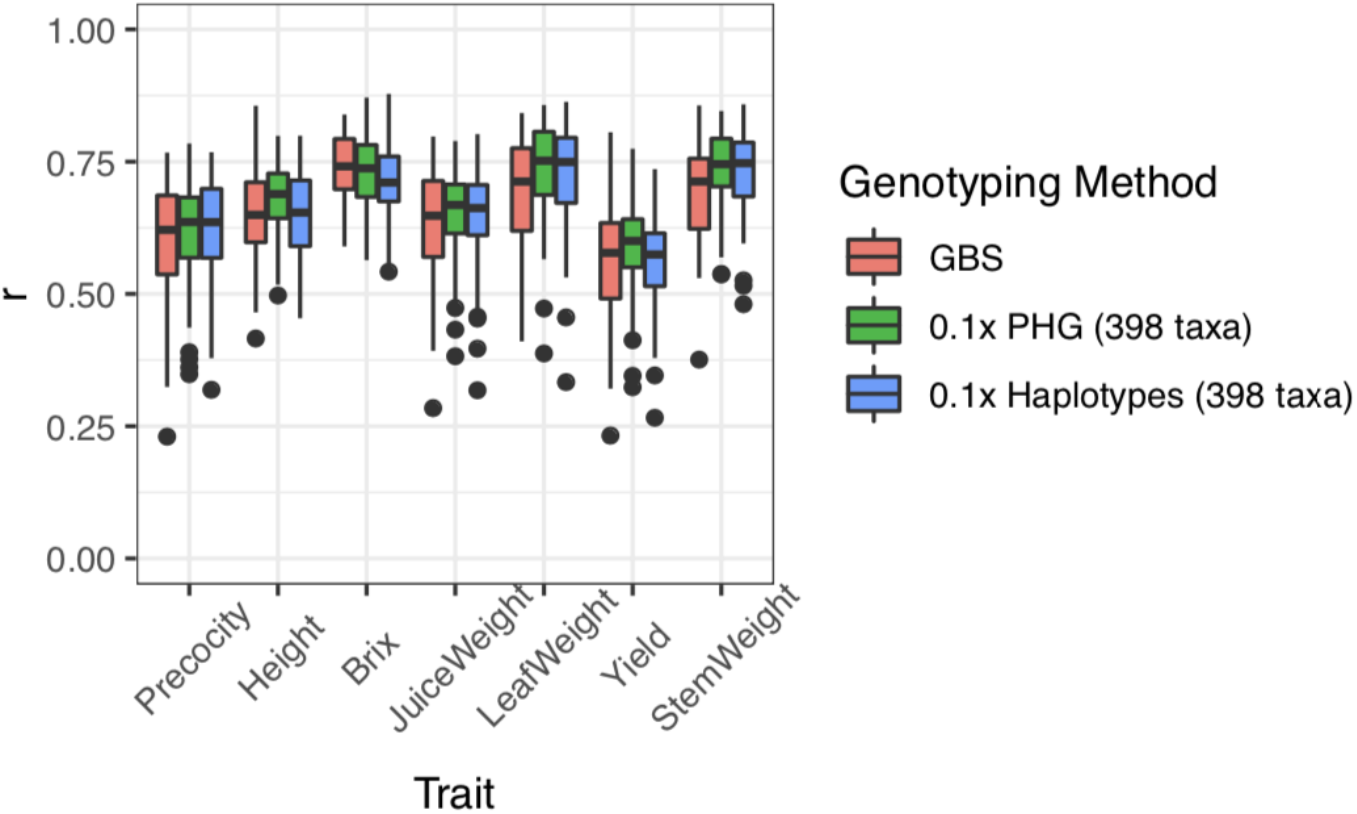
Genomic prediction accuracies from diversity PHG SNPs (green) or haplotypes (blue) match prediction accuracies of GBS (pink).

## Notes

#### Summary of Updates

Author names updated

https://bitbucket.org/bucklerlab/p_sorghumphg/src/master/

https://doi.org/10.25739/nccb-0k78

## References

Alonso-Blanco, C., J. Andrade, C. Becker, F. Bemm, J. Bergelson, et al. 2016. 1,135 Genomes Reveal the Global Pattern of Polymorphism in Arabidopsis thaliana. Cell 166(2): 481–491. doi: 10.1016/j.cell.2016.05.063.

Altschul, S.F., T.L. Madden, A.A. Schäffer, J. Zhang, Z. Zhang, et al. 1997. Gapped BLAST and PSI-BLAST: A new generation of protein database search programs. Nucleic Acids Res. 25(17): 3389–3402. doi: 10.1093/nar/25.17.3389.

Anderson, C.B., B.K. Franzmayr, S.W. Hong, A.C. Larking, T.C. Stijn, et al. 2018. Protocol: A versatile, inexpensive, high-throughput plant genomic DNA extraction method suitable for genotyping-by-sequencing. Plant Methods 14(1): 1–10. doi: 10.1186/s13007-018-0336-1.

Bates, D., M. Mächler, B.M. Bolker, and S.C. Walker. 2015. Fitting linear mixed-effects models using lme4. J. Stat. Softw. 67(1). doi: 10.18637/jss.v067.i01.

Baym, M., S. Kryazhimskiy, T.D. Lieberman, H. Chung, M.M. Desai, et al. 2015. Inexpensive Multiplexed Library Preparation for Megabase-Sized Genomes. PLoS One 10(5): 1–15. doi: 10.1371/journal.pone.0128036.

Bennetzen, J.L., J. Schmutz, H. Wang, R. Percifield, J. Hawkins, et al. 2012. Reference genome sequence of the model plant Setaria. Nat. Biotechnol. 30(6): 555–561. doi: 10.1038/nbt.2196.

Bernardo, R., and J. Yu. 2007. Prospects for genomewide selection for quantitative traits in maize. Crop Sci. 47(3): 1082–1090. doi: 10.2135/cropsci2006.11.0690.

Bouchet, S., M.O. Olatoye, S.R. Marla, R. Perumal, T. Tesso, et al. 2017. Increased Power To Dissect Adaptive Traits in Global Sorghum Diversity Using a Nested Association Mapping Population. Genetics 206(June): 573–585. doi: 10.1534/genetics.116.198499/-/DC1.1.

Bradbury, P.J., Z. Zhang, D.E. Kroon, T.M. Casstevens, Y. Ramdoss, et al. 2007. TASSEL: Software for association mapping of complex traits in diverse samples. Bioinformatics 23(19): 2633–2635. doi: 10.1093/bioinformatics/btm308.

Browning, B.L., and S.R. Browning. 2016. Genotype Imputation with Millions of Reference Samples. Am. J. Hum. Genet. 98(1): 116–126. doi: 10.1016/j.ajhg.2015.11.020.

Bukowski, R., X. Guo, Y. Lu, C. Zou, B. He, et al. 2018. Construction of the third-generation Zea mays haplotype map. Gigascience 7(4): 1–12. doi: 10.1093/gigascience/gix134.

Elshire, R.J., J.C. Glaubitz, Q. Sun, J.A. Poland, K. Kawamoto, et al. 2011. A robust, simple genotyping-by-sequencing (GBS) approach for high diversity species. PLoS One 6(5): 1–10. doi: 10.1371/journal.pone.0019379.

Endelman, J.B. 2011. Ridge Regression and Other Kernels for Genomic Selection with R Package rrBLUP. Plant Genome J. 4(3): 250. doi: 10.3835/plantgenome2011.08.0024.

Fresnedo-Ramírez, J., S. Yang, Q. Sun, A. Karn, B.I. Reisch, et al. 2019. Computational analysis of ampseq data for targeted, high-throughput genotyping of amplicons. Front. Plant Sci. 10(May): 1–8. doi: 10.3389/fpls.2019.00599.

Girma, G., H. Nida, A. Seyoum, M. Mekonen, A. Nega, et al. 2019. A large-scale genome-wide association analyses of ethiopian sorghum landrace collection reveal loci associated with important traits. Front. Plant Sci. 10(May): 1–15. doi: 10.3389/fpls.2019.00691.

Heffner, E.L., A.J. Lorenz, J.L. Jannink, and M.E. Sorrells. 2010. Plant breeding with Genomic selection: Gain per unit time and cost. Crop Sci. 50(5): 1681–1690. doi: 10.2135/cropsci2009.11.0662.

Heslot, N., J. Jannink, and M.E. Sorrells. 2015. Perspectives for Genomic Selection Applications and Research in Plants. (February): 1–12. doi: 10.2135/cropsci2014.03.0249.

Jiang, Y., R.H. Schmidt, and J.C. Reif. 2018. Haplotype-Based Genome-Wide Prediction Models Exploit Local Epistatic Interactions Among Markers. G3&#58; Genes|Genomes|Genetics 8(5): 1687–1699. doi: 10.1534/g3.117.300548.

Li, H. 2013. seqtk: Toolkit for processing sequences in FASTA/Q formats.: https://github.com/lh3/seqtk.

Li, H., and R. Durbin. 2009. Fast and accurate short read alignment with Burrows-Wheeler transform. Bioinformatics 25(14): 1754–1760. doi: 10.1093/bioinformatics/btp324.

Li, H., B. Handsaker, A. Wysoker, T. Fennell, J. Ruan, et al. 2009. The Sequence Alignment/Map format and SAMtools. Bioinformatics 25(16): 2078–2079. doi: 10.1093/bioinformatics/btp352.

Lozano, R., Gazave, E., dos Santos, J. P. R., Stetter, M., Valluru, R., Bandillo, N., Fernandes, S. B., Brown, P. J., Shakoor, N., Mockler, T., Ross-Ibarra, J., Buckler, E. S., Gore, M. A. 2019. Comparative evolutionary analysis and prediction of deleterious mutation patterns between sorghum and maize. Biorxiv.

Mace, E.S., S. Tai, E.K. Gilding, Y. Li, P.J. Prentis, et al. 2013. Whole-genome sequencing reveals untapped genetic potential in Africa’s indigenous cereal crop sorghum. Nat. Commun. 4: 2320. doi: 10.1038/ncomms3320.

Mace, E.S., J.F. Rami, S. Bouchet, P.E. Klein, R.R. Klein, et al. 2009. A consensus genetic map of sorghum that integrates multiple component maps and high-throughput Diversity Array Technology (DArT) markers. BMC Plant Biol. 9: 1–14. doi: 10.1186/1471-2229-9-13.

McCormick, R.F., S.K. Truong, A. Sreedasyam, J. Jenkins, S. Shu, et al. 2017. The Sorghum bicolor reference genome: improved assembly, gene annotations, a transcriptome atlas, and signatures of genome organization. Plant J. doi: 10.1111/tpj.13781.

Meuwissen, T.H.E., B.J. Hayes, and M.E. Goddard. 2001. Prediction of total genetic value using genome-wide dense marker maps. Genetics 157(4): 1819–1829. doi: 11290733.

Morris, G.P., P. Ramu, S.P. Deshpande, C.T. Hash, T. Shah, et al. 2013. Population genomic and genome-wide association studies of agroclimatic traits in sorghum. Proc. Natl. Acad. Sci. 110(2): 453–8. doi: 10.1073/pnas.1215985110.

Muleta, K.T., G. Pressoir, and G.P. Morris. 2019a. Optimizing genomic selection for a sorghum breeding program in Haiti: a simulation study. Genes|Genomes|Genetics (February): 1–22. doi: 10.1534/g3.118.200932.

Muleta, K.T, Winans, N., Felderhoff, T., Charles, J. R., Pressoir, G., Armstrong, J. S., Morris, G. P. 2019b. Recent evolutionary rescue of sorghum in the Americas required sixty years of global germplasm exchange. Biorxiv.

Ouyang, S., W. Zhu, J. Hamilton, H. Lin, M. Campbell, et al. 2007. The TIGR Rice Genome Annotation Resource: Improvements and new features. Nucleic Acids Res. 35(SUPPL. 1): 8–11. doi: 10.1093/nar/gkl976.

Pereira, M.G., and M. Lee. 1995. Identification of genomic regions affecting plant height in sorghum and maize. Theor. Appl. Genet. 90: 380–388. doi: 10.1007/BF00221980.

Poland, J., J. Endelman, J. Dawson, J. Rutkoski, S. Wu, et al. 2012. Genomic Selection in Wheat Breeding using Genotyping-by-Sequencing. Plant Genome J. 5(3): 103. doi: 10.3835/plantgenome2012.06.0006.

Rabiner, L.R. 1989. A tutorial on HMM and selected Applications in Speech Recognition. Proc. IEEE 77(2): 257–286.

Ramstein, G.P., S.E. Jensen, and E.S. Buckler. 2018. Breaking the curse of dimensionality to identify causal variants in Breeding 4. Theor. Appl. Genet. (0123456789). doi: 10.1007/s00122-018-3267-3.

Ribaut, J.M., M.C. de Vicente, and X. Delannay. 2010. Molecular breeding in developing countries: challenges and perspectives. Curr. Opin. Plant Biol. 13(2): 213–218. doi: 10.1016/j.pbi.2009.12.011.

Schnable, P.S., D. Ware, R.S. Fulton, J.C. Stein, F. Wei, et al. 2009. The B73 Maize Genome: Complexity, Diversity, and Dynamics. Science (80-.). 326(June): 1112–1116.

Sentieon DNASeq, “Sentieon,” 2018. [Online]. Available: https://www.sentieon.com/products/

Shakoor, N., R. Nair, O. Crasta, G. Morris, A. Feltus, et al. 2014. A Sorghum bicolor expression atlas reveals dynamic genotype-specific expression profiles for vegetative tissues of grain, sweet and bioenergy sorghums. BMC Plant Biol. 14(1): 35. doi: 10.1186/1471-2229-14-35.

Valluru, R., E.E. Gazave, S.B. Fernandes, J.N. Ferguson, R. Lozano, et al. 2019. Deleterious Mutation Burden and Its Association with Complex Traits in Sorghum (Sorghum bicolor). Genetics 211(3): 1075 LP – 1087. doi: 10.1534/genetics.118.301742.

Vogel, J.P., D.F. Garvin, T.C. Mockler, J. Schmutz, D. Rokhsar, et al. 2010. Genome sequencing and analysis of the model grass Brachypodium distachyon. Nature 463(7282): 763–768. doi: 10.1038/nature08747.

Watson, A., S. Ghosh, M.J. Williams, W.S. Cuddy, J. Simmonds, et al. 2018. Speed breeding is a powerful tool to accelerate crop research and breeding. Nat. Plants 4(1): 23–29. doi: 10.1038/s41477-017-0083-8.

Zheng, X., D. Levine, J. Shen, S.M. Gogarten, C. Laurie, et al. 2012. A high-performance computing toolset for relatedness and principal component analysis of SNP data. Bioinformatics 28(24): 3326–3328. doi: 10.1093/bioinformatics/bts606.

Zou, Y., M.G. Mason, Y. Wang, E. Wee, C. Turni, et al. 2017. Nucleic acid purification from plants, animals and microbes in under 30 seconds. PLOS Biol. 15(11): e2003916. doi: 10.1371/journal.pbio.2003916.

